# Benchmarking single-cell foundation models in a zero-shot setting

**DOI:** 10.64898/2026.08.03.739553

**Authors:** Yasmine Gaballa, Somaia Ahmed, Tamim Abdelaal

**Affiliations:** Systems and Biomedical Engineering Department, Faculty of Engineering Cairo University, 12613 Giza, Egypt

## Abstract

Single-cell foundation models have recently emerged as a promising approach for learning general-purpose representations from large-scale transcriptomic data. These models are trained on millions of cells and are designed to transfer their learned representations to a wide range of downstream tasks. However, their practical benefits compared to traditional approaches are still not fully understood. This study evaluates four foundation models, namely scGPT, SCimilarity, UCE, and Transcriptformer, across four downstream tasks: cell type annotation, human data integration, cross-species data integration, and protein expression prediction. Embeddings generated by each model were assessed using multiple public single-cell datasets and compared against conventional machine learning baselines. Performance was measured using task-specific evaluation metrics, including classification, integration, and regression metrics. The results showed that foundation model embeddings did not consistently outperform traditional approaches. In the cell type annotation task, baseline methods achieved the strongest performance across most datasets. For protein expression prediction, however, embeddings from the foundation models generally produced more accurate predictions than the baseline, with SCimilarity achieving the lowest prediction error and Transcriptformer obtaining the highest correlation scores. In the data integration task, all foundation models produced moderate results, while scVI (the baseline) achieved the strongest integration performance. Overall, the results suggest that current single-cell foundation models provide useful representations for some downstream tasks in zero-shot conditions but do not yet offer a universal replacement for task-specific methods. Their effectiveness remains dependent on the application and evaluation setting.

## 1. Introduction

Recent advances in genomics have greatly improved our understanding of the molecular processes that govern cellular function. By studying genes and their activity, researchers can gain insights into biological processes that support applications such as disease diagnosis, drug discovery, and precision medicine. As genomic technologies continue to evolve, the volume and complexity of biological data have grown substantially, creating new opportunities and challenges for data-driven analysis.

One of the most influential developments in this field is single-cell RNA sequencing (scRNA-seq), which enables gene expression to be measured at the level of individual cells. Unlike traditional bulk sequencing approaches that average signals across large cell populations, scRNA-seq captures cellular heterogeneity and reveals distinct cell types and states within complex tissues [1, 2]. The resulting data are typically represented as a high-dimensional matrix, where rows correspond to cells, columns correspond to genes, and each entry represents a gene’s expression level in a particular cell. Single-cell data support a wide range of downstream analytical tasks, including cell type annotation, disease classification, domain integration, expression and perturbation prediction. However, these tasks are often complicated by challenges such as data sparsity, high dimensionality, technical noise, and batch effects. These limitations have motivated the development of machine learning approaches capable of learning meaningful representations from large-scale genomic datasets [3].

In recent years, foundation models have emerged as a powerful paradigm for representation learning. Originally developed for natural language processing and computer vision, these models are pretrained on vast amounts of data and can be adapted to multiple downstream tasks. Their ability to capture general patterns and transferable knowledge has inspired their application to biological data [4]. Single-cell foundation models, such as UCE, extend these principles to large-scale gene expression datasets. By learning biologically meaningful representations during pretraining, these models aim to support a variety of downstream analyses. As a result, they have become a promising direction for advancing computational genomics and improving our understanding of complex biological systems [5–10].

Several foundation models have been developed for single-cell genomics. Although these models all aim to learn useful information from large-scale biological data, they differ in how they are trained and in the tasks they are designed to support. Geneformer was one of the first transformer-based foundation models developed for single-cell genomics. It was trained on millions of single-cell transcriptomes and learns relationships between genes and cellular states. The model has been used in several applications, including cell state analysis, gene network studies, and perturbation-related research [5]. scGPT is a foundation model developed for single-cell and multi-omics data analysis. It applies generative pretraining to gene expression data in a way that is inspired by large language models. The goal is to learn cell representations that can be reused across different biological tasks. The model has shown promising results in applications such as cell type annotation, data integration, and perturbation analysis [6]. Universal Cell Embeddings (UCE) was proposed to create cell embeddings that can be used across different datasets, tissues, and species. The model was trained on a large collection of single-cell data from various species and aims to capture common biological patterns that remain useful in different settings. This makes UCE a suitable candidate for tasks involving data integration and knowledge transfer [7]. SCimilarity was developed to identify cells with similar biological characteristics. The model learns embeddings that place similar cells close to each other, making it easier to compare cells across different datasets. This approach can support tasks such as cell annotation, cell retrieval, and large-scale exploration of cell atlases [8]. Transcriptformer is a foundation model trained on single-cell datasets collected from many different species. By learning from data spanning a large species range, the model aims to capture biological patterns that are shared across organisms. This makes it particularly relevant for tasks that involve transferring information between species [9]. Nicheformer extends the idea of foundation models to spatial omics data. In addition to gene expression information, it also considers the spatial location of cells and their surrounding cellular environment. This allows the model to study cell interactions and tissue organization in greater detail [10]. scBERT is a transformer-based foundation model developed for single-cell analysis. Unlike many earlier approaches, the model adapts the BERT architecture to gene expression data and learns contextual relationships between genes through masked prediction during pretraining. This allows the model to capture complex gene-gene interactions and transfer the learned representations to downstream tasks such as cell type annotation [11]. scFoundation is a foundation model trained on a very large corpus of single-cell data and designed for large-scale single-cell transcriptomic analysis. The model learns general representations from diverse gene expression datasets and aims to support a wide range of downstream applications, including cell type annotation, perturbation prediction, and gene expression analysis [12].

Although all of these models are designed to learn transferable representations from biological data, they focus on different aspects of cellular biology. For example, scGPT and Geneformer focus on large-scale transcriptomic pretraining; UCE and Transcriptformer emphasize generalization across datasets and species; and Nicheformer incorporates spatial information. Because of such differences, model performance may vary depending on the task being evaluated.

As single-cell foundation models have become more widely used, researchers have started to compare their performance across different biological tasks. These studies provide useful insights into the current models, but they also show that benchmarking remains a challenging problem. *Kedzierska et al.* evaluated two single-cell foundation models (scGPT and Geneformer) in zero-shot settings, where they were tested in batch integration without fine-tuning. Their results showed that strong pretrained representations do not always lead to the best downstream performance and highlight the importance of careful model evaluation [13]. *Csendes et al.* focused on benchmarking two foundation models (scGPT and scFoundation) for post-perturbation RNA sequencing prediction. Their work examined how well pretrained models can predict cellular responses after biological perturbations. While the study provided valuable results for this specific application, it was limited to a single downstream task [14]. *Boiarsky et al.* argued that current foundation models (such as scBERT and scGPT, which were tested in the study) require more thorough evaluation before broad conclusions can be made about their capabilities. The study showed that model performance, specifically in the task of cell type annotation, can depend strongly on the selected datasets, evaluation metrics, and experimental design, emphasizing the need for more consistent benchmarking practices [15].

Reviewing the existing benchmarking studies showed that most evaluations focus on a limited number of models or downstream tasks. Many studies compare only one or two foundation models, while others focus on a single application such as cell type annotation or perturbation prediction [13,14,15]. In addition, tasks such as protein expression prediction and cross-species integration are rarely included in benchmarking frameworks despite their biological relevance. As a result, it remains difficult to assess how foundation models perform across a broader range of single-cell analysis tasks.

To address this gap, four downstream tasks were selected. Cell type annotation (both standalone and as a label transfer follow-up experiment after data integration) was included because it is one of the most common and fundamental tasks in single-cell analysis. Human data integration was used to assess the ability of a model to combine datasets from different studies, laboratories, or disease conditions while preserving biological structure. Cross-species integration was included to evaluate whether biological information can be transferred between different organisms. Finally, protein expression prediction was selected because it assesses whether the learned representations capture biological information beyond cell identity and can be used to estimate protein abundance from gene expression profiles. Based on these tasks, four single-cell foundation models were selected for evaluation: scGPT, UCE, SCimilarity, and TranscriptFormer. These models were chosen because they represent different approaches of representation learning in single-cell analysis and collectively support the range of tasks considered in this study. In particular, UCE and Transcriptformer were trained on data from multiple species, making them especially relevant for cross-species integration experiments. Together, the selected models provide a diverse set of architectures and training strategies for evaluating the strengths and limitations of current single-cell foundation models. By evaluating all models using the same datasets, metrics, and experimental procedures, this work aims to provide a fair and consistent comparison across all tested downstream tasks.

## 2. Methods

### 2.1. Overview of Evaluation Pipeline

This study evaluates four single-cell foundation models using a unified benchmarking pipeline based on pretrained embeddings. The models are not fine-tuned on any downstream task. Instead, embeddings are extracted directly from pretrained checkpoints and used as fixed representations across all experiments.

The evaluation is carried out on four downstream tasks: cell type annotation, human data integration, cross-species integration, and protein expression prediction. Each task tests a different aspect of model performance, including classification ability, robustness to batch effects, transfer across species, and prediction of protein levels. In total, nine single-cell datasets are used in the evaluation. These datasets vary in size and biological context. Some datasets are reused across multiple tasks to ensure consistent and comparable evaluation settings.

The overall pipeline is shown in Figure 1. A dataset is first passed through a pretrained model to extract cell embeddings. These embeddings are then used as input for task-specific models, which are evaluated using standard metrics. Overall, this approach allows a fair comparison of models by focusing on the quality of learned representations rather than differences in training or fine-tuning.

**Figure 1:**
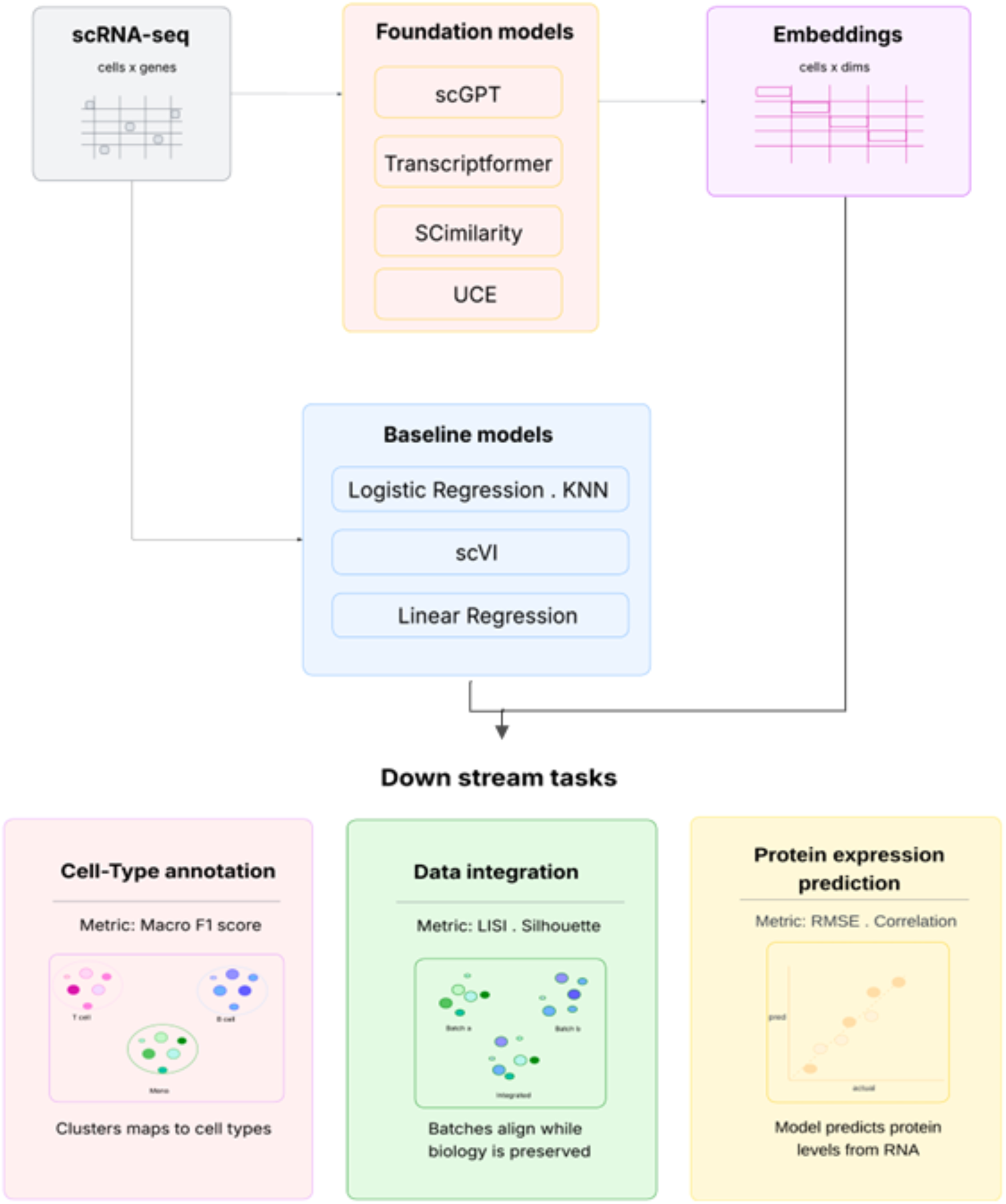
The overall evaluation pipeline

### 2.2. Tested Models

In this work, 4 models were tested: scGPT, Transcriptformer, UCE, and SCimilarity. Table 1 provides an overview of these models, summarizing their architectures, pre-training data, and embedding dimensions. The following subsections discuss each model individually.

**Table 1:**
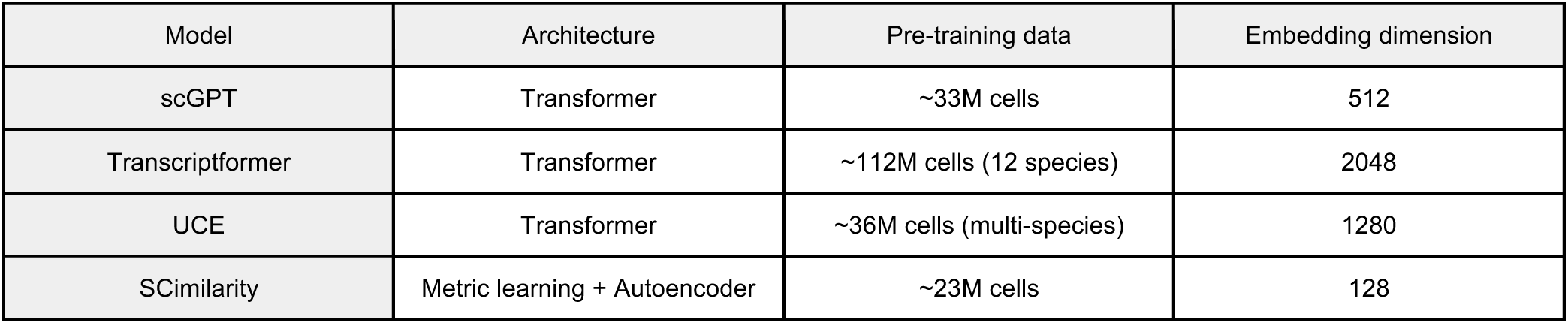
Summary of the tested models and their details.

| Model | Architecture | Pre-training data | Embedding dimension |
| --- | --- | --- | --- |
| scGPT | Transformer | ~33M cells | 512 |
| Transcriptformer | Transformer | ~112M cells (12 species) | 2048 |
| UCE | Transformer | ~36M cells (multi-species) | 1280 |
| SCimilarity | Metric learning + Autoencoder | ~23M cells | 128 |

#### 2.2.1. scGPT

scGPT [5] is a Transformer-based foundation model for single-cell analysis. It extends the Generative Pre-trained Transformer (GPT) paradigm to gene expression data. Specifically, genes are treated as tokens, and gene relationships are learned during pretraining. The Transformer utilizes a multi-head self-attention mechanism that learns dependencies between genes. This allows the model to learn interactions between genes and complex expression patterns across each cell. Embeddings extracted from the model can be applied to a variety of downstream tasks such as cell type annotation, data integration, and perturbation prediction. The model was pretrained on a collection of publicly available single-cell RNA sequencing datasets containing more than 33 million human cells [16] from different tissues and biological conditions [6]. Training on such a large and diverse dataset allows the model to learn general patterns in gene expression that can be transferred to new datasets. The input to scGPT is a single-cell gene expression matrix in AnnData [17] format. Genes are matched to the model vocabulary before being processed by the model. Each cell is then converted into a fixed-length embedding vector that summarizes its gene expression profile. The output is an embedding matrix of size N x 512 containing one 512-dimensional embedding for each cell in the dataset, and these embeddings are used directly in all downstream tasks.

#### 2.2.2. Transcriptformer

TranscriptFormer [8] is a foundation model for single-cell analysis based on the Transformer architecture. Similar to other transformer-based models, it uses self-attention layers to learn relationships between genes and capture patterns within gene expression data. The model follows a generative learning approach, where it learns representations directly from gene expression profiles and produces embeddings that can be transferred to downstream tasks. Unlike many existing foundation models, TranscriptFormer was designed for cross-species analysis and aims to learn biological patterns that are shared across different organisms. The model was trained on a large cross-species collection of single-cell transcriptomic data containing up to 112 million cells from 12 species. Training on data from multiple species allows the model to learn more general representations that can be applied across different biological systems [9]. The input to TranscriptFormer is a single-cell gene expression matrix in AnnData format. Gene identifiers are mapped to the model vocabulary before being processed by the model. Each cell is then converted into a fixed-length embedding vector representing its gene expression profile. The output is an embedding matrix of size N x 2048 containing one 2048-dimensional embedding for each cell in the dataset, and these embeddings are used directly in all downstream evaluation tasks.

#### 2.2.3. SCimilarity

SCimilarity [8] is a cell representation autoencoder-based model that combines supervised metric learning with unsupervised reconstruction learning. The model is trained using two objectives: a triplet loss that pulls cells with matching cell-type labels closer together in the embedding space and a mean squared error (MSE) reconstruction loss that preserves information from the original gene expression profiles. The training corpus includes 412 studies covering approximately 23.4 million cells. From this, 56 studies (about 7.9 million cells) are used for training, and 15 studies (about 1.4 million cells) are used for testing [8]. The remaining data consists of unannotated or unused studies. The input to SCimilarity is a single-cell gene expression matrix with gene symbols in AnnData format. Before embedding generation, genes are aligned to the model’s reference gene set and preprocessing steps are performed through the model’s built-in pipeline. Each cell is then encoded into a fixed-length representation. The output is a 128-dimensional embedding vector for every cell, resulting in an embedding matrix of size *N* × 128 for a dataset containing *N* cells. These embeddings are used directly in all downstream tasks.

#### 2.2.4. UCE

Universal Cell Embeddings (UCE) [7] is a transformer-based foundation model designed to learn cell representations that generalize across datasets and species. The original UCE model consists of a 33- layer transformer architecture and was pretrained on around 36 million single cells collected from around 300 studies. The training corpus includes data across dozens of tissues and eight species, allowing the model to capture biological patterns shared across different organisms. In this study, embeddings were generated using the UCE-100M checkpoint through the Hugging Face Accelerator [18]. The input to UCE is a single-cell gene expression profile with gene symbols. Each cell is processed independently to produce a high-dimensional representation of it. For every input cell, the model generates a 1280- dimensional embedding vector, resulting in an embedding matrix of size *N* × 1280 for a dataset containing *N* cells. These embeddings are extracted using the pretrained model and later used in all downstream tasks.

### 2.3. Datasets

A total of nine datasets were used in this study. Seven datasets consist exclusively of human cells, one dataset (Han et al. [19]) is from macaque samples, and one (Baron et al. [20]) contains both human and mouse cells. The datasets were distributed across four tasks, with some datasets sometimes being used in more than one task. Table 2 provides a summary of the datasets, including their tissue of origin, species, number of cells, and the tasks for which they were used to evaluate. Before evaluation, all datasets were inspected to ensure compatibility with the selected models and downstream analysis pipelines. Minor dataset-specific adjustments, such as standardizing metadata fields and gene identifiers, were performed when required.

**Table 2:**
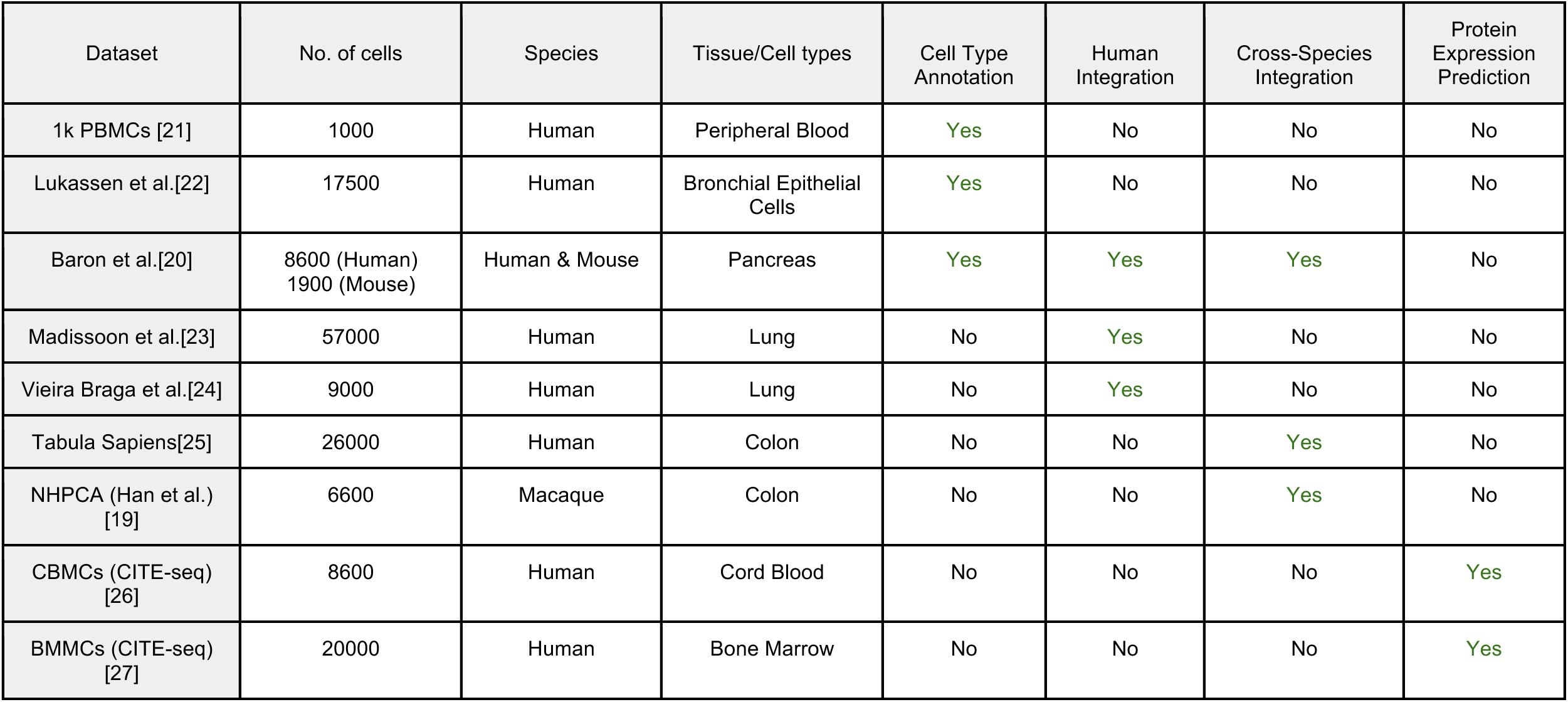
Description of the datasets used in the study and the tasks in which they were used for evaluation.

For the baseline experiments, gene expression data were normalized, and highly variable genes (HVGs) were selected to reduce dimensionality and retain the most informative features. These processed features were then used as input for the classification and regression models. In addition, principal component analysis (PCA) was applied in visualization tasks when needed to improve the clarity of the resulting plots.

### 2.4. Downstream tasks

#### 2.4.1. Cell Type Annotation

Cell type annotation evaluates the ability of a model to distinguish between different cell populations. The objective is to assign the correct cell type label to each cell. Accurate cell type annotation indicates that the extracted embeddings preserve biologically meaningful information that can be used to separate different cell clusters. To evaluate this task, embeddings generated by each foundation model were used as input features for two classifiers: Logistic Regression and K-Nearest Neighbors (KNN). The same classifiers were also applied directly to the original gene expression after normalization and highly variable genes (HVG) selection to establish baseline performance. For the KNN classifier, the number of neighbors was fixed (k=15) across all experiments. For this specific task, 5-fold cross-validation was used to preserve class distribution. The predicted output for each cell is a cell type label corresponding to one of the classes present in the dataset.

#### 2.4.2. Data Integration

The data integration task was used to evaluate whether the embeddings generated by each foundation model could align cells from different datasets while preserving biologically meaningful cell-type structure. Two integration settings were considered: human-only integration and cross-species integration. The human-only experiments assessed the ability of the models to reduce dataset-specific differences between human datasets, while the cross-species experiments evaluated whether cells with similar biological identities could be grouped together despite originating from different species. For each experiment, embeddings generated by the foundation models were combined and analyzed in a shared embedding space. To further assess the quality of the integration, label transfer was performed using the same classification pipeline described in the cell type annotation task. A classifier was trained on the source dataset and then used to predict cell-type labels in the target dataset. If biologically similar cells were successfully aligned after integration, the transferred labels were expected to match the ground-truth annotations of the target cells. For the baseline integration method, scVI [28] was used. It was trained on the combined datasets after normalization, logarithmic transformation, and selection of the top 4,000 highly variable genes. The scVI architecture was configured with two hidden layers and a latent dimension of 30. After training, the learned latent representations were extracted and used as the integrated embeddings for downstream analysis, same as the four foundation models. The output of this task is a shared embedding space containing cells from multiple datasets.

#### 2.4.3. Protein Expression Prediction

Protein expression prediction examines how well the learned representations capture the relationship between gene expression and protein abundance. In this task, protein levels are estimated from the embeddings generated by each foundation model. CITE-seq data is used for evaluation, where gene and protein expressions are measured simultaneously at the single-cell level [26,27]. Good performance suggests that the embeddings retain biologically relevant information that extends beyond basic cell-type differences. For evaluation, a Linear Regression model was trained with a 5-fold cross-validation using the embeddings produced by each foundation model. The same regression model was also applied directly to the original gene expression after the same preprocessing done on the datasets from the former tasks to provide a baseline for comparison. The predicted output consists of continuous protein expression values for each cell.

### 2.5. Evaluation Metrics

Depending on the downstream task at hand, some quantitative metrics were used along with some visualization techniques in order to interpret the results. Classification metrics were used for cell type annotation, integration metrics were used for human and cross-species data integration, and regression metrics were used for protein expression prediction.

For the cell type annotation task, performance was evaluated using Accuracy, Precision, and Macro F1- score. Accuracy measures the proportion of correctly classified cells among all cells in the dataset 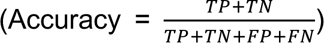. The values range from 0 to 1, where 1 indicates perfect classification performance and 0 indicates completely incorrect predictions. Precision measures the proportion of correctly predicted cell labels among all predictions assigned to a given class 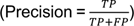. Values range from 0 to 1, with higher values indicating fewer false positive predictions and more reliable cell type classifications. The F1-score combines precision and recall into a single metric (F1 = 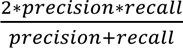, where Recall = 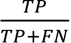). Macro F1-score is calculated by computing the F1-score independently for each class and then averaging across all classes. Values range from 0 to 1. Higher values indicate that the model performs consistently across both common and rare cell types, making this metric particularly useful when class distributions are imbalanced.

For human and cross-species data integration, performance was evaluated using the Silhouette Score and the Local Inverse Simpson’s Index (LISI). The Silhouette Score 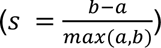 measures how well cells are grouped within clusters while remaining separated from other clusters. Values range from -1 to 1. A value close to 1 indicates well-separated clusters, a value near 0 indicates overlapping clusters, and negative values suggest incorrect clustering. For Cell Type Silhouette scores, higher values indicate better preservation of biological structure, while for Dataset Silhouette scores, lower values are generally preferred because successful integration should reduce separation between datasets. LISI [29] evaluates the diversity of neighboring cells in the embedding space and is commonly used to assess integration quality. Higher dataset LISI values indicate better mixing between datasets, while lower cell-type LISI values indicate stronger preservation of biological structure.

For protein expression prediction, performance was evaluated using Pearson correlation and Root Mean Squared Error (RMSE). Pearson correlation 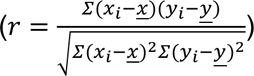 measures the strength of the linear relationship between predicted (y) and observed (x) protein expression values. The values range from -1 to 1. Values closer to 1 indicate a stronger positive relationship and therefore better predictions. RMSE is the square root of MSE and provides an error measure in the same units as the original protein expression values 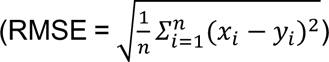. Values range from 0 to positive infinity, and lower values indicate better predictions.

## 3. Results

### 3.1. Cell type annotation: No improvement over baseline

Cell type annotation is a crucial downstream task in single-cell analysis, in which cells are assigned biological identities based on their gene expression profiles. Since single-cell foundation models are pretrained on large-scale transcriptomic data, they are hypothesised to encode biologically meaningful representations that could improve cell-type discrimination beyond conventional features representations. This experiment assesses whether embeddings from such models enhance supervised annotation performance relative to a standard highly variable gene (HVG) baseline.

Cell type annotation was performed using Logistic Regression and K-Nearest Neighbors classifiers trained on embeddings extracted from four foundation models: UCE, SCimilarity, scGPT, and Transcriptformer. These embeddings were compared against the baseline. Performance was evaluated using Accuracy, Precision, and Macro F1-score, with primary emphasis on Macro F1-score due to class imbalance. Experiments were conducted using four datasets: PBMCs, HBECs, Human Pancreas (Baron et al.), and Mouse Pancreas.

For the PBMC dataset (Figure 2A), the baseline achieved strong and stable performance across metrics. UCE showed comparable results to the baseline, while SCimilarity and scGPT underperformed, particularly in Macro F1-score when paired with Logistic Regression. Transcriptformer achieved the best performance among foundation models and slightly exceeded the baseline; however, the improvement remained marginal. Overall, these results indicate that while foundation model embeddings preserve biologically relevant structure, they yield limited gains when cell populations are already well-separated.

**Figure 2:**
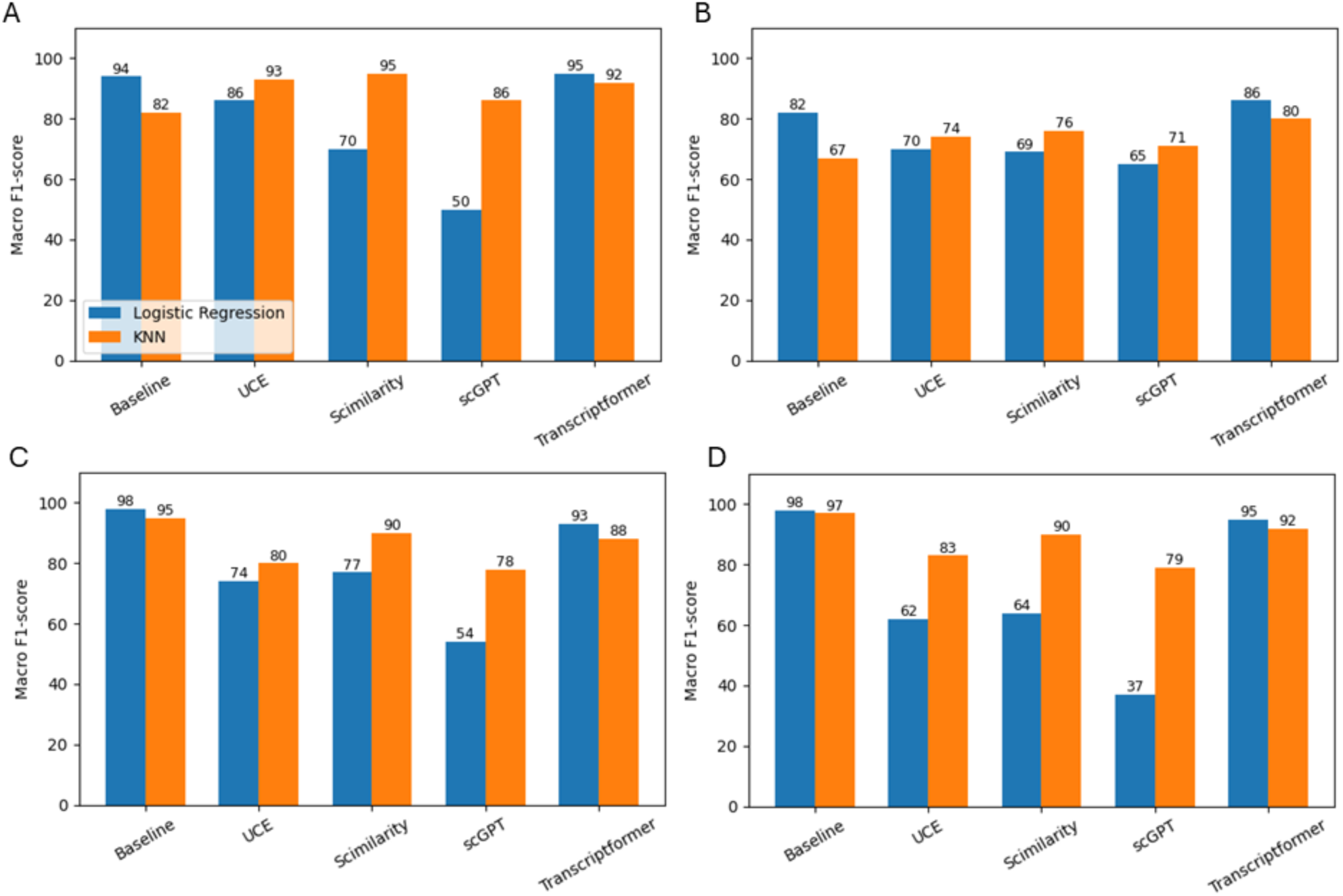
Cell type annotation performance using HVG features and single-cell foundation model embeddings across four benchmark datasets. **(A)** PBMC dataset. **(B)** HBEC dataset. **(C)** Human Pancreas dataset. **(D)** Mouse Pancreas dataset. Performance is reported using Macro F1- score for Logistic Regression and K-Nearest Neighbors classifiers. Across all datasets, the HVG baseline remains highly competitive and often matches or exceeds the performance of foundation-model embeddings. Although Transcriptformer generally achieves the strongest results among the foundation models, no embedding method consistently outperforms the baseline, indicating the limited benefit of current foundation- model representations for supervised cell type annotation.

Consistent trends were observed across the HBEC and Pancreatic datasets (Figures 2B–D). Transcriptformer remained the most competitive embedding-based model, achieving the highest Macro F1-score among foundation models in most cases. However, performance differences between embeddings and the baseline remained small, with the baseline often matching or outperforming most models. This pattern was consistent across both human (HBEC, Baron Pancreas) and mouse pancreas datasets, indicating that HVG features already capture sufficient discriminatory information for cell type classification in these settings.

Overall, the pretrained foundation model embeddings retain biologically meaningful information but do not consistently improve cell type annotation performance over HVG-based representations. Across all datasets and classifiers, no foundation model systematically outperformed the baseline, reinforcing the conclusion that cell type annotation shows no improvement over baseline when using current foundation model embeddings.

### 3.2. Data Integration: Limited Advantages over Specialized Integration Models

#### 3.2.1. Human

Batch integration aims to remove unwanted technical and cohort-associated variation introduced by differences in sequencing protocols, laboratories, sample preparation, and sequencing machines, while preserving underlying biological structure, particularly cell type identity. Successful integration, therefore, requires balancing two competing objectives: cells originating from different batches should become well mixed, while cells belonging to the same biological cell type should remain clearly separated.

To evaluate this trade-off, various datasets containing batch-associated technical variation were analyzed using the foundation models to generate a shared batch-corrected embedding space in comparison to scVI as a baseline model. The integration performance was assessed using both Silhouette and LISI metrics. For Silhouette scores, desirable embeddings exhibit high cell type Silhouette values together with low Dataset Silhouette values, indicating strong biological separation and effective batch mixing. For LISI, lower Cell Type LISI values indicate tighter cell type clustering, whereas higher Dataset LISI values indicate greater batch diversity within local neighborhoods. UMAP[30] projections were also generated for each model and dataset to visually assess dataset mixing and cell type separation.

As shown in Figure 3A, the Human Pancreas dataset illustrates the general integration behavior observed across the human datasets. The original unintegrated data exhibited strong apparent cell type structure (Cell Type Silhouette ≈ 0.25) but limited batch mixing (Dataset Silhouette ≈ 0.016, Dataset LISI ≈ 1.11), indicating that batch effects contributed to the observed clustering. Following integration, clear differences emerged between methods. SCimilarity achieved the most favorable overall balance, combining the highest Cell Type Silhouette (≈ 0.26) with the lowest (best) Dataset Silhouette (≈ -0.04), indicating strong preservation of biological structure alongside effective batch correction. The LISI analysis showed that SCimilarity maintained cell-type compactness comparable to scVI, although scVI achieved stronger batch mixing according to Dataset LISI. UCE achieved moderate performance, preserving reasonable cell-type structure while partially correcting batch effects. In contrast, scGPT achieved stronger batch mixing, particularly according to Dataset LISI, but this was accompanied by reduced cell-type separation. Transcriptformer showed poor cell-type preservation and failed to achieve strong batch correction, resulting in the weakest overall integration performance. These quantitative findings are further supported by the UMAP visualizations in Figure 4. The original unintegrated embedding exhibits visible separation between healthy and diseased samples, whereas SCimilarity and UCE preserve compact cell-type clusters while improving condition mixing. In contrast, Transcriptformer produces a diffuse and overlapping embedding with limited cell-type separation, consistent with its low Cell Type Silhouette score.

**Figure 3:**
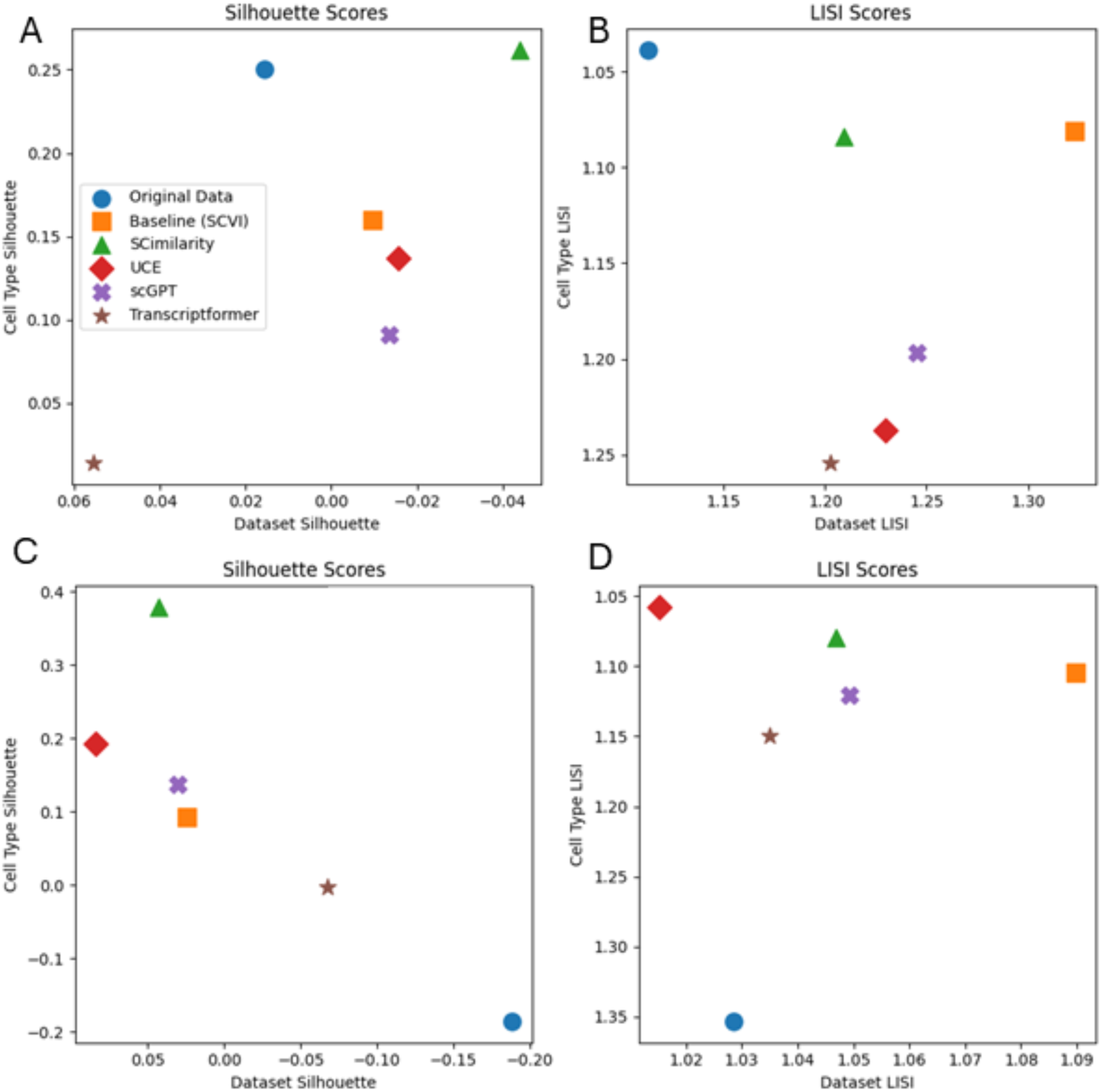
Batch integration performance on the Human Pancreas (A) and Human Lung (B) datasets. Scatter plots show Silhouette scores (left) and LISI scores (right) for the original unintegrated data, the SCVI baseline, and four foundation models (UCE, SCimilarity, scGPT, and TranscriptFormer). For the Silhouette plots, the best performance lies in the upper-right region, corresponding to high Cell Type Silhouette and low Dataset Silhouette values (x-axis reversed), indicating strong preservation of biological structure and effective batch mixing. For the LISI plots, the best performance also occupies the upper-right region, corresponding to low Cell Type LISI (y-axis reversed) and high Dataset LISI values. Each point represents the aggregate integration result of a single method. The strongly negative silhouette scores observed for the original Lung data highlight the severe batch-induced distortion present before integration.

**Figure 4:**
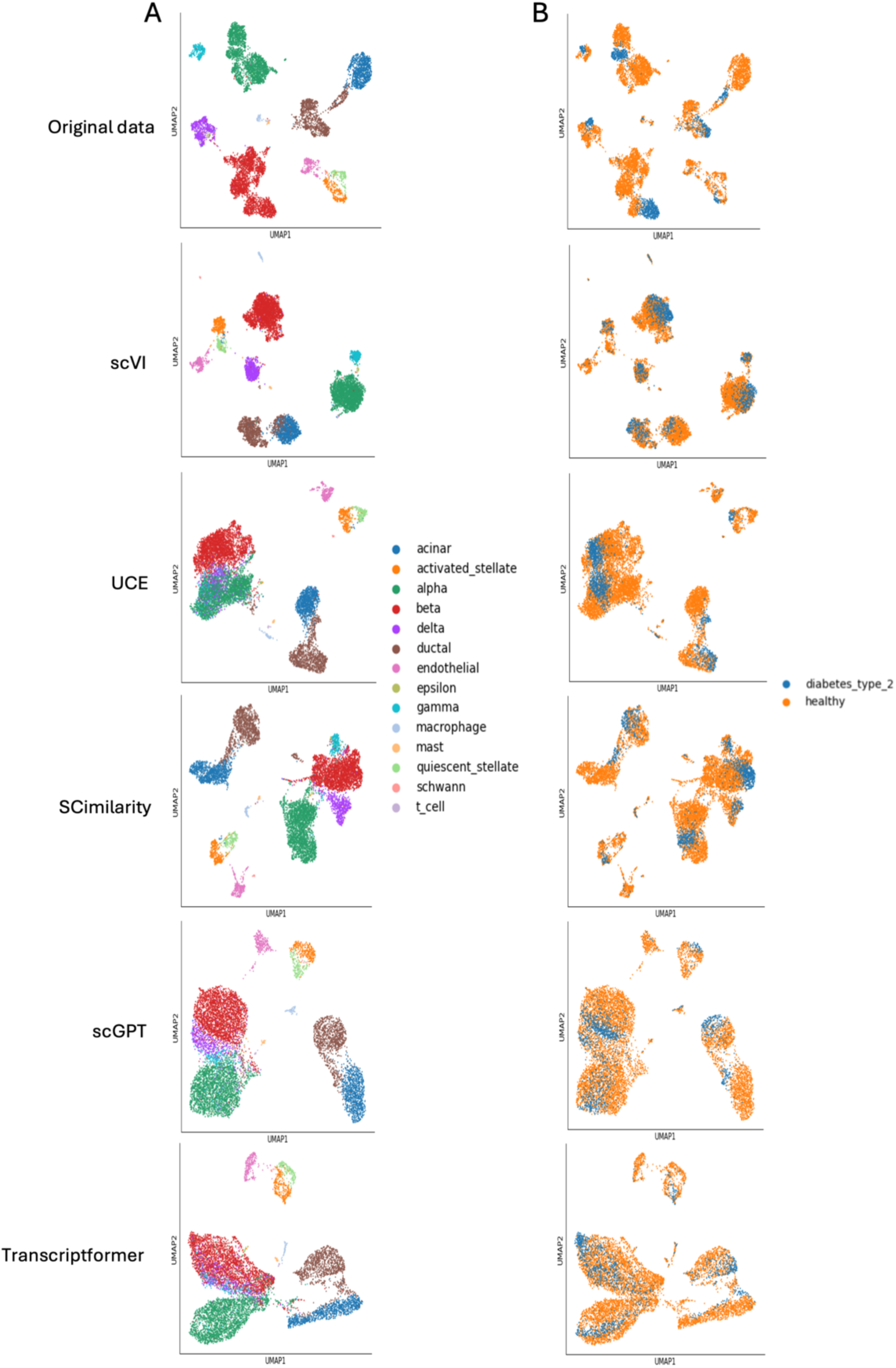
UMAP visualizations of batch integration on the Human Pancreas dataset. Columns correspond to Original data (no integration), scVI, UCE, SCimilarity, scGPT, and Transcriptformer. The left column (A) is colored by cell type, while the right column (B) is colored by condition (type 2 diabetes vs. healthy), serving as a proxy for batch identity. An ideal integration preserves compact, well-separated cell type clusters while mixing cells from both conditions within each cluster. The original data exhibit clear cell type structure but noticeable condition separation. scVI achieves the strongest condition mixing but with reduced cell type compactness. UCE and SCimilarity maintain distinct cell type clusters while providing moderate batch mixing. scGPT produces broader, less compact clusters, whereas Transcriptformer yields a largely continuous embedding with substantial cluster overlap, consistent with its low cell type silhouette score (Figure 3A).

A similar pattern was observed on the Lung dataset (Figure 3B), although the integration task was considerably more challenging. The original data showed strongly negative Cell Type and Dataset Silhouette scores, indicating severe batch-induced distortion in the raw feature space. SCimilarity achieved the strongest cell-type separation (Cell Type Silhouette ≈ 0.38) but retained residual batch effects. UCE also preserved biological structure, achieving a relatively high Cell Type Silhouette (≈0.192) and the lowest Cell Type LISI; however, it exhibited the weakest batch correction, with the highest Dataset Silhouette (≈0.084) and lowest Dataset LISI. In contrast, scVI achieved the strongest batch mixing according to Dataset LISI while maintaining moderate cell-type separation, representing the most balanced overall integration performance on this dataset. scGPT occupied an intermediate position, achieving better batch mixing than UCE and SCimilarity but without reaching the integration performance of scVI. Transcriptformer achieved the lowest Dataset Silhouette, suggesting strong batch mixing according to this metric; however, this was accompanied by near-complete loss of cell-type separation and poor Cell Type LISI, indicating over-correction of biological signals.

Overall, the integration results reveal that the utility of foundation model embeddings for batch correction is highly dataset dependent and does not consistently surpass the specialized scVI baseline. On the Pancreas dataset, SCimilarity achieved the best silhouette-based balance between cell-type preservation and batch mixing, although scVI provided the strongest batch mixing according to LISI. On the more challenging Lung dataset, foundation models often exhibited a trade-off between preserving biological structure and removing batch effects. SCimilarity achieved the strongest cell-type separation but retained residual batch structure, whereas UCE preserved cell identity but showed the weakest batch mixing. In contrast, scVI achieved the most consistent balance between batch correction and biological preservation across the evaluated datasets, highlighting the advantage of methods specifically optimized for integration. scGPT generally occupied an intermediate position, offering moderate batch correction but sacrificing some cell-type resolution. Transcriptformer generally overcorrected the data, leading to poor preservation of biological structure.

#### 3.2.2. Cross-species

Cross-species integration presents a more challenging setting than within-species batch correction. In this case, the goal is not only to reduce technical variation but also to align transcriptomic representations across evolutionarily distinct organisms. This introduces a fundamental tension: while shared cell type programs should be aligned, species-specific biological differences must not be fully collapsed. As a result, successful integration requires preserving meaningful biological divergence while still enabling partial correspondence between equivalent cell types across species.

The human–mouse pancreas dataset was used to evaluate cross-species integration between evolutionarily distant organisms. The original unintegrated data showed a cell type silhouette of approximately 0.08 and a Dataset Silhouette of approximately 0.23, reflecting strong species-driven separation in the raw feature space (Figure 5A). Among the evaluated methods, SCimilarity achieved the strongest cell-type separation according to Silhouette scores (Cell Type Silhouette ≈ 0.17) while reducing species structure (Dataset Silhouette ≈ 0.07). In contrast, scVI achieved slightly lower cell-type separation (≈0.13) but provided the strongest species mixing according to both Dataset Silhouette (≈ 0.02) and Dataset LISI, while also achieving the lowest Cell Type LISI. This highlights how different evaluation metrics emphasize distinct aspects of cross-species integration, with Silhouette scores favoring biological separation and LISI emphasizing local species mixing. scGPT achieved moderate species mixing but showed weaker cell-type compactness, while UCE showed limited improvement in cross-species alignment and retained substantial species separation. Transcriptformer largely failed to preserve biological structure, exhibiting near-zero Cell Type Silhouette. Overall, the results illustrate the trade-off between aligning shared cellular programs and preserving species-specific biological variation.

**Figure 5:**
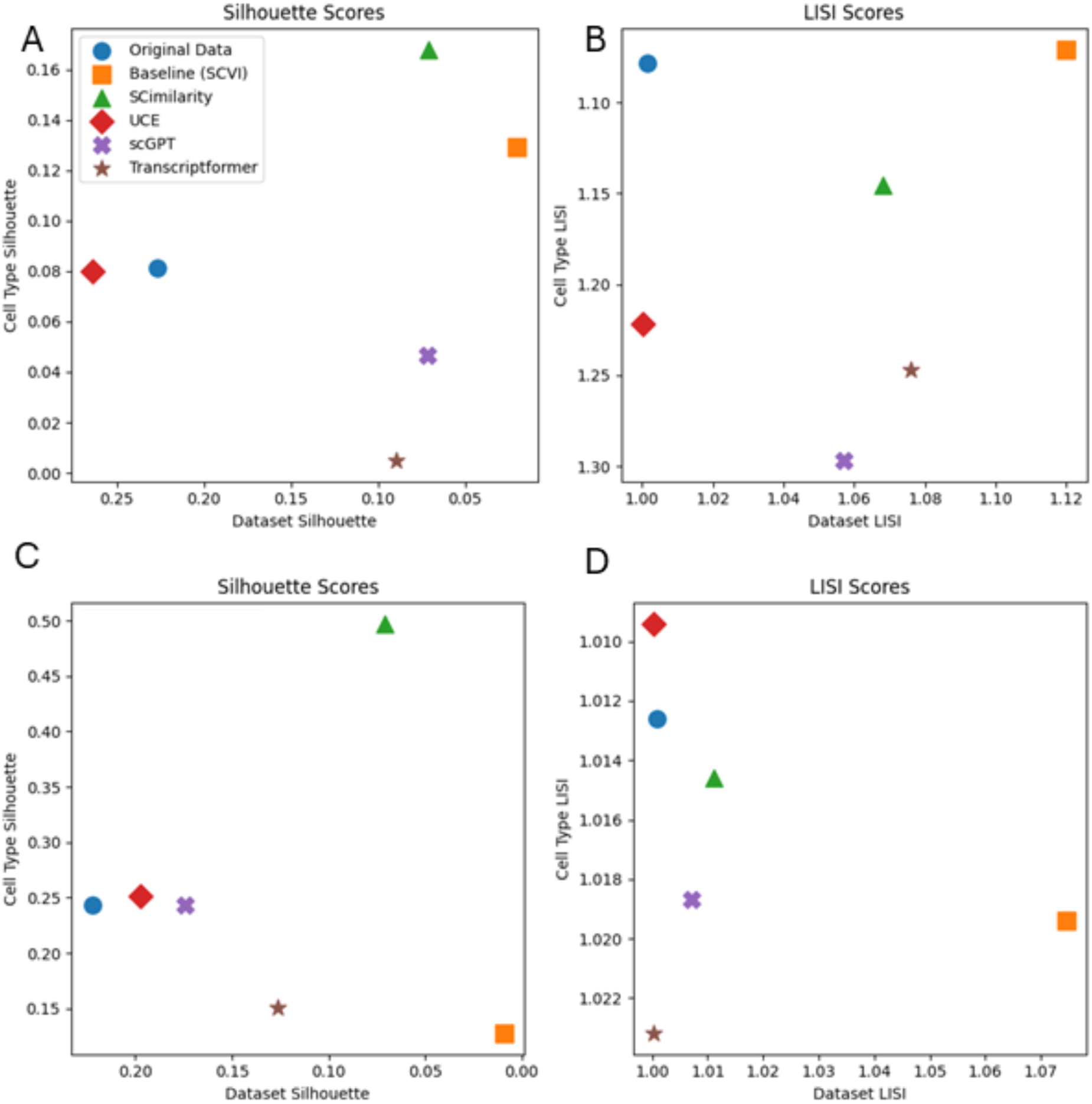
Batch integration performance on the cross-species datasets, including the Pancreas (human–mouse) (A) and Colon (human– monkey) (B). Scatter plots show Silhouette scores (left) and LISI scores (right) for the original unintegrated data, the SCVI baseline, and four single-cell foundation models (UCE, SCimilarity, scGPT, and TranscriptFormer). In both settings, the batch covariate corresponds to species identity, making these cross-species integration tasks. For the silhouette plots, the best performance lies in the upper-right region, indicating high Cell Type Silhouette and low Dataset Silhouette values (x-axis reversed), reflecting strong biological structure preservation and effective species-level mixing. For the LISI plots, the best performance also corresponds to low Cell Type LISI (y-axis reversed) and high Dataset LISI values in the upper-right region. In the cross-species Pancreas dataset, all methods retain some degree of species-level separation, reflecting the challenge of aligning shared cellular programs while preserving genuine evolutionary differences. In the Colon dataset, SCimilarity achieves the highest Cell Type Silhouette (∼0.50) while reducing species separation, whereas scVI achieves the strongest species mixing (Dataset Silhouette near zero and highest Dataset LISI) at the cost of reduced biological separation. Overall, the narrow performance range across LISI scores highlights the difficulty of balancing species alignment with preservation of species-specific biological variation.

The human–monkey colon dataset represents a less extreme cross-species scenario. The original data already showed relatively strong cell type structure (Cell Type Silhouette ≈ 0.25) and substantial species separation (Dataset Silhouette ≈ 0.22) (Figure 5B). SCimilarity achieved the strongest cell-type preservation (≈ 0.50) while still reducing species separation (≈ 0.07). The LISI analysis showed that SCimilarity maintained Cell Type LISI values comparable to UCE, indicating compact biological clustering. In contrast, scVI achieved the strongest species mixing according to both Dataset Silhouette and Dataset LISI but at the expense of reduced cell-type separation, suggesting partial overcorrection. UCE and scGPT preserved moderate biological structure, with scGPT achieving slightly better species mixing than UCE, but neither reached the balanced performance observed for SCimilarity or the stronger alignment achieved by scVI. Transcriptformer again showed reduced cell-type preservation and only moderate improvements in species mixing.

Across both cross-species datasets, integration performance showed a clear trade-off between species mixing and preservation of biological identity. SCimilarity achieved the strongest cell-type separation according to Silhouette scores, particularly in the human–monkey colon dataset, whereas scVI consistently achieved stronger species mixing and was favored by LISI-based evaluation. UCE and scGPT showed moderate or limited improvements depending on the dataset, while Transcriptformer consistently failed to preserve meaningful biological structure. These findings indicate that cross-species integration remains challenging, as successful methods must balance alignment of conserved cellular programs with preservation of genuine evolutionary differences.

#### 3.2.3. Label Transfer

Label transfer evaluates whether cell type information learned from one dataset can be successfully transferred to another dataset. Unlike standard cell type annotation, where training and testing occur within the same dataset, label transfer is performed after integration and therefore assesses whether the integrated embedding aligns biologically similar cells across batches or species. Successful transfer requires cells belonging to the same cell type to occupy similar regions of the embedding space regardless of their dataset or species of origin while maintaining sufficient separation from other cell types.

Performance was evaluated using Accuracy, Precision, and Macro F1-score for Logistic Regression (LR) and KNN classifiers, with Macro F1-score serving as the primary evaluation metric due to its robustness to class imbalance. Results are presented separately for within-species and cross-species transfer settings in Figure 6.

**Figure 6:**
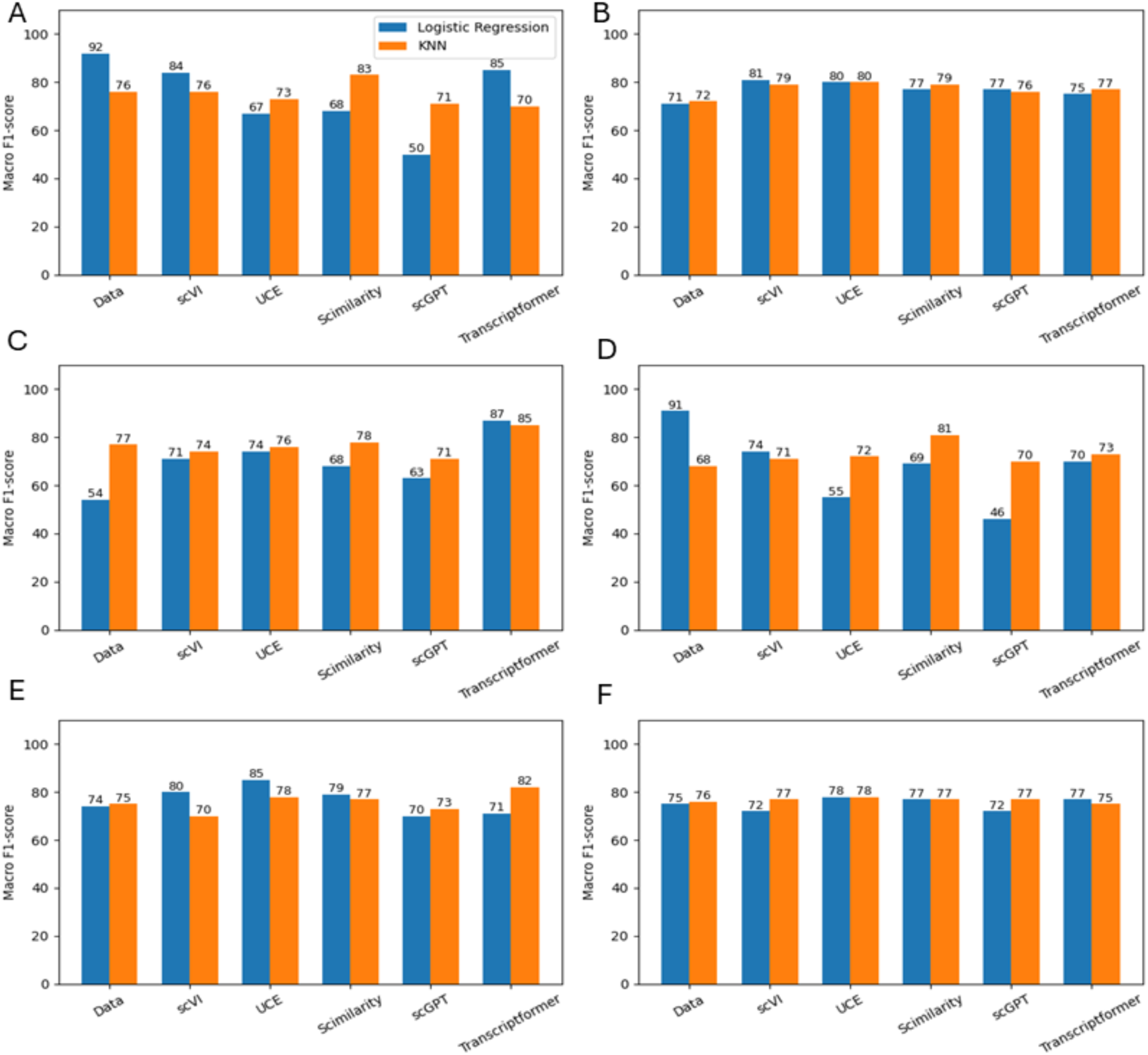
Label transfer performance for within-species and cross-species experiments. **Figures 6a–6b** show within-species label transfer on the Baron Pancreas and Lung datasets, respectively, while **Figures 6c–6f** show cross-species transfer on the Baron Pancreas (Human→Mouse and Mouse→Human) and Colon (Human→Monkey and Monkey→Human) datasets. Bar charts reportMacro F1-score for Logistic Regression (LR) and KNN classifiers trained on embeddings from the raw data, scVI, UCE, SCimilarity, scGPT, and Transcriptformer. Within-species experiments evaluate the transfer of labels across batches from the same species, whereas cross-species experiments assess the ability of embeddings to generalize cell type information across evolutionary boundaries. Across the within-species datasets, performance differences between methods are generally modest, with the raw data baseline remaining highly competitive. In contrast, the cross-species experiments reveal larger performance differences between representations, highlighting the increased difficulty of transferring cell type information across species. Transcriptformer achieves the strongest performance in the Human→Mouse pancreas setting, while UCE and SCimilarity provide the most consistent results across the colon transfer tasks. Overall, the results illustrate how embedding quality influences the preservation and transferability of cell type structure under increasingly challenging biological shifts.

Within-species transfer generally produced strong performance across all methods. On the Baron Pancreas dataset (Figure 6A), the raw data baseline achieved the highest overall performance, reaching Macro F1-scores of 92% and 76% under Logistic Regression and KNN, respectively. Among the foundation models, SCimilarity and Transcriptformer produced the strongest KNN results, while scGPT showed notably weaker Logistic Regression performance. In contrast, performance differences were considerably smaller on the Lung dataset (Figure 6B), where all methods achieved similar Macro F1- scores ranging between approximately 71% and 81%. These results suggest that within-species transfer remains largely accessible using conventional gene-expression features, limiting the advantage provided by learned embeddings.

Cross-species transfer provides a more demanding evaluation because cell type representations must remain consistent despite evolutionary divergence between species. As shown in Figures 6C and 6D, transfer between human and mouse pancreatic cells proved substantially more challenging than within-species transfer. In the Human→Mouse (train → test) direction, Transcriptformer achieved the strongest performance overall, reaching Macro F1-scores of 87% and 85% under Logistic Regression and KNN, respectively. However, performance declined noticeably in the reverse Mouse→Human direction, suggesting asymmetry in the alignment between human and mouse transcriptomic representations. SCimilarity achieved the strongest KNN performance in this setting, while scGPT recorded its lowest scores across all transfer experiments.

Figures 6E and 6F present the corresponding results for human–monkey colon transfer. Compared with the pancreas experiments, overall performance was higher and more stable, likely reflecting the closer evolutionary relationship between the two species. UCE and SCimilarity consistently achieved the strongest results, while Transcriptformer produced the highest KNN performance in the Human→Monkey direction. Unlike the pancreas datasets, differences between methods were relatively small in the Monkey→Human task.

Overall, label transfer results demonstrate that foundation model embeddings provide limited benefit for within-species transfer, where the raw data baseline often remains competitive or superior. In contrast, larger differences emerged in cross-species settings, where learned representations were required to capture cell type relationships that generalize across evolutionary boundaries. UCE and SCimilarity offered the most consistent performance across datasets, while Transcriptformer showed strong but highly dataset-dependent behavior. These findings suggest that the primary advantage of single-cell foundation models lies not in improving conventional within-species annotation, but in supporting transfer learning scenarios where biological variation between training and testing datasets is substantial.

### 3.3. Protein Expression Prediction: Substantial Improvement over Baseline Predictions

Protein expression prediction is one of the downstream tasks in single-cell analysis, where the objective is to predict protein expression levels from gene expression profiles. Since single-cell foundation models are pretrained on large-scale transcriptomic data, they are expected to learn representations that better capture regulatory signals relevant to protein expression compared to conventional gene-expression features. This experiment evaluates whether embeddings from these models improve supervised protein prediction performance relative to a standard highly variable gene (HVG) baseline.

Protein expression prediction was evaluated on the CBMC and Bone Marrow CITE-seq datasets, which provide paired measurements of gene expression and surface protein abundance at the single-cell level. In this task, only the transcriptomic (gene expression) profiles were provided as input to the foundation models to generate cell embeddings. These embeddings were then used as features for a linear regression model trained to predict the corresponding protein expression levels. The performance of the resulting predictions was evaluated using two complementary metrics: Pearson correlation, which measures the agreement between predicted and observed protein levels (higher values indicate better performance), and RMSE, which measures the prediction error (lower values indicate better performance). Embeddings from four foundation models, UCE, SCimilarity, scGPT, and Transcriptformer, were compared against a baseline using highly variable gene (HVG) expression features.

On the CBMCs dataset (Figures 7A and 7B), all foundation models consistently outperformed the HVG baseline across both metrics. The baseline achieved a median Pearson correlation of approximately 0.612 and a median RMSE of about 0.438, with noticeable variability across proteins. In contrast, all embedding-based models improved predictive performance and reduced variability. scGPT achieved the highest median Pearson correlation (≈0.76), followed closely by UCE and SCimilarity (≈0.744– 0.749), while Transcriptformer also performed strongly (≈0.728). In terms of RMSE, scGPT and SCimilarity achieved the lowest errors (≈0.32–0.328), corresponding to an approximate 25% reduction compared to the baseline. Additionally, all foundation models showed narrower performance distributions, indicating more stable predictions across proteins.

**Figure 7:**
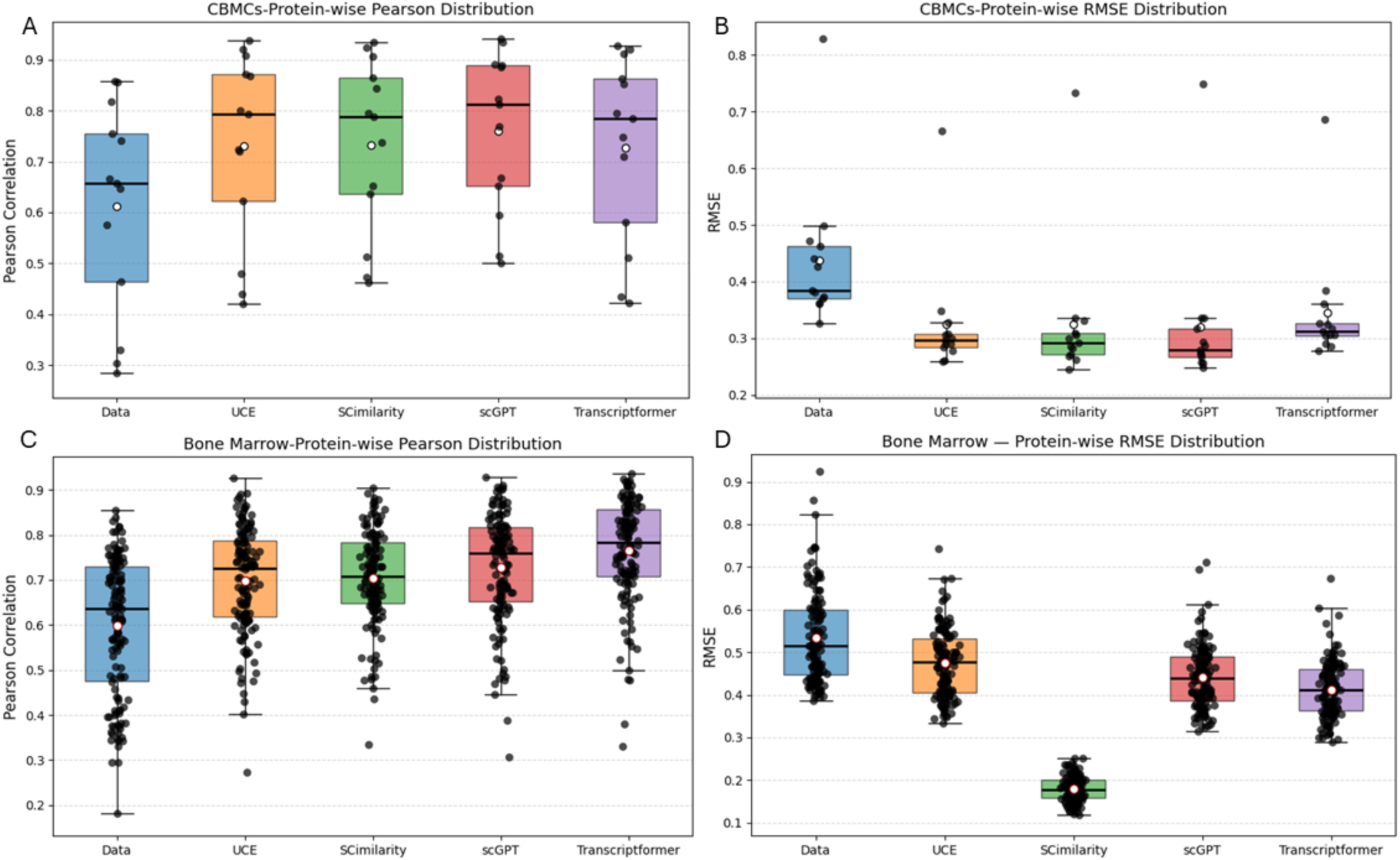
Protein expression prediction performance on the CBMC and Bone Marrow CITE-seq datasets. Figures7A–7B show results on the CBMC dataset, while Figures 7C–7D show results on the Bone Marrow dataset. Box plots report the distribution of protein-wise Pearson correlation coefficients (left) and RMSE values (right) obtained using linear regression models trained on embeddings from the HVG baseline, UCE, SCimilarity, scGPT, and Transcriptformer. Across both datasets, all foundation models consistently outperform the HVG baseline, demonstrating that pretrained representations encode biologically meaningful signals beyond conventional gene expression features. scGPT achieves the highest median correlation on the CBMC dataset, while Transcriptformer achieves the highest correlation on Bone Marrow. For RMSE, SCimilarity and scGPT achieve the lowest prediction errors on CBMC, whereas SCimilarity provides the largest reduction in prediction error on Bone Marrow.

Figures 7C and 7D present the corresponding results for the Bone Marrow dataset. Similar trends were observed, although score distributions were generally wider than those in CBMCs. The HVG baseline achieved a median Pearson correlation of approximately 0.575 and a median RMSE of approximately 0.536. All foundation models improved performance relative to the baseline. Transcriptformer achieved the highest median Pearson correlation (≈0.768), followed by scGPT (≈0.728). For RMSE, SCimilarity achieved the lowest median value (≈0.181), corresponding to a reduction of more than 65% relative to the baseline. While SCimilarity achieved the smallest prediction error, Transcriptformer obtained the highest correlation, indicating that the model with the lowest absolute error is not necessarily the model with the strongest agreement in protein-level variation.

Overall, all foundation model embeddings substantially improved protein expression prediction compared with HVG-based features. The improvements were consistent across datasets and evaluation metrics, although the best-performing model depended on the dataset and metric considered. scGPT achieved the strongest overall performance on CBMCs, while Transcriptformer achieved the highest correlation, and SCimilarity achieved the lowest RMSE on Bone Marrow. These findings suggest that pretrained single-cell foundation models capture biologically meaningful signals that are highly beneficial for protein expression prediction.

## 4. Discussion

The results across cell type annotation, data integration, label transfer, and protein expression prediction reveal that the utility of single-cell foundation models is highly task-dependent. While foundation model embeddings provided limited benefits for conventional cell type annotation, they showed greater value in integration, transfer learning, and protein prediction tasks that require more generalizable biological representations. This suggests that the primary strength of current foundation models lies not in improving standard classification performance but in capturing broader biological structure that can support more complex downstream analyses.

For cell type annotation, the limited improvement over the HVG baseline is not entirely surprising and is consistent with recent benchmarking studies. In our experiments, classifiers were trained and evaluated within the same dataset, a setting in which highly variable genes already provide strong discriminatory power because major cell populations are often well separated in expression space. Under these conditions, foundation model embeddings may offer little additional benefit, as the supervised classifier can learn dataset-specific decision boundaries directly from the available features. Similar observations have been reported by Kedzierska et al. (2025) [13], showing that scGPT and Geneformer do not consistently outperform simpler HVG-based and established single-cell analysis methods across cell-type-related tasks, and by Steiner et al. (2025) [31], concluding that the performance of foundation models for cell type classification remains highly dependent on both the dataset and the model architecture. Together, these findings suggest that current foundation models preserve biologically meaningful information but do not fundamentally improve class separability when training and evaluation occur within the same dataset. However, a different pattern emerged in the integration and label transfer experiments, where the ability to learn transferable biological representations becomes more important than maximizing classification performance within a single dataset.

The integration and label transfer results provide additional insight into the strengths and limitations of foundation model representations. Interestingly, UCE did not consistently emerge as the strongest performer in cross-species integration despite being designed to learn universal cellular representations across species. Instead, SCimilarity frequently achieved better preservation of cell type structure, while UCE generally provided a more balanced trade-off between biological preservation and batch mixing. One possible explanation is that cross-species integration requires not only aligning shared biological programs but also preserving genuine species-specific differences, making complete alignment inherently difficult. More broadly, it is important to note that the scVI baseline was trained specifically for each dataset and task, allowing it to learn dataset-specific representations optimized for integration. In contrast, the foundation models were evaluated using their pretrained embeddings in a zero-shot manner without task-specific fine-tuning. The competitive performance achieved under this setting is therefore encouraging and suggests that further gains may be possible through supervised adaptation or fine-tuning. This interpretation is further supported by the label transfer results, where foundation models showed their greatest advantage in cross-species settings. The consistency between the integration and label transfer experiments suggests that models that better preserve biologically meaningful structure after integration are also more effective at transferring cell identities across datasets and species.

Protein expression prediction was the task that benefited most consistently from foundation model representations. Across both datasets, all foundation models substantially outperformed the HVG baseline, improving both Pearson correlation and RMSE. Unlike cell type annotation, where highly variable genes already contain sufficient information to separate major cell populations, protein expression prediction requires modeling complex and often non-linear relationships between transcriptomic and proteomic measurements. The superior performance of embedding-based models therefore suggests that large-scale pretraining enables the capture of latent biological signals that extend beyond the information contained in individual highly variable genes. The particularly strong performance observed in the Bone Marrow dataset, especially for SCimilarity, further indicates that certain representations may encode regulatory patterns that are more closely aligned with protein-level variation. Although improvements in RMSE and Pearson correlation were not always proportional, this is expected because the two metrics evaluate different aspects of prediction quality: Pearson correlation measures the consistency of predicted expression trends, whereas RMSE measures absolute prediction error. Overall, these findings suggest that the greatest value of current single-cell foundation models lies in tasks that require rich biological representation learning, where pretrained embeddings can capture complex molecular relationships that are difficult to recover from conventional gene-expression features alone.

It is important to note that we relied in our evaluation of the foundation models on pretrained embeddings, without any fine-tuning. While fine-tuning may enhance performance on some tasks, it was omitted to ensure a fair and consistent comparison across all models.Furthermore, some models may require different preprocessing steps or require specific formats for the input data, like variations in gene identifiers (gene symbols compared to Ensembl IDs) and different minimum number of features requirements. Not all datasets are applied to every model, which reduces the diversity in the datasets used. Lastly, we focused on four downstream tasks, however, evaluating additional downstream tasks, like perturbation prediction or trajectory inference, would allow a wider assessment of the foundation models capabilities.

## 5. Conclusion

The main aim of this study is to assess whether single-cell foundation models would provide better representations than traditional approaches across various downstream tasks. Overall, our results showed that foundation models can be beneficial, however, this is task dependent; being most effective when the downstream task required capturing more complex biological relationships, while simpler approaches remained highly competitive for cell type annotation.

## Code Availability

The code to reproduce the results is available in the GitHub repository, at https://github.com/somaiaahmed/Benchmarking-Single-Cell-Foundation-Models-in-Zero-Shot-Setting

## Notes

### Competing Interest Statement

The authors have declared no competing interest.

